# Human Brain Mapping of Homotopic Functional Affinity

**DOI:** 10.1101/2024.01.09.574929

**Authors:** Li-Zhen Chen, Xi-Nian Zuo

## Abstract

Homotopic positions are defined as the two areas with opposite but equal horizonal coordinates in the standard symmetric brain space. Characterizing similarity between two homotopic areas, brain homotopy represents a typical feature of the two hemispheres for both structure and function. Functional homotopy provides important perspectives for understanding neural correlates of cognition and behavior. Despite the decisive role of spatial geometric constraints and homophilic attachment on the human connectome, traditional practices in mapping functional homotopy only considered the temporal correlations of functional timeseries between homotopic areas, but ignored the homophily factors in generative connectivity models. Here, we proposed a novel method for functional homotopy analysis, namely Homotopic Functional Affinity (HFA). This method quantifies the homotopic affinity as the cosine distance of the full-brain functional connectivity profiles or fingerprints between the homotopic areas. HFA captures both geometric constraints (homotopic location) and homophily (affinity) simultaneously. By leveraging the resting-state fMRI data from the Human Connectome Project (HCP) and the Chinese HCP (CHCP), we mapped the 700ms-2mm high spatiotemporal resolution HFA and evaluated its test-retest reliability with linear mixed models, exhibiting generally fair-to-substantial reliable measurements of individual differences in HFA. The lowest HFA observed in the temporo-parietal junction (TPJ) inspired to perform an edge-detection algorithm on its surface render and derived three clearly differentiable and adjacent TPJ subregions: the anterior TPJ (TPJa), the central TPJ (TPJc), and the posterior TPJ (TPJp). We further validated the HFA for the three TPJ regions through a set of comprehensive analyses, including the delineation of their functional connectivity fingerprints, the meta-analysis of their cognitive functions, and their task-activation correlation. Finally, we linked the cortical HFA map to those multimodal brain maps of gene expression, evolution, myelination, functional hierarchy, and cognitive association. The systematic subregion analysis revealed the complex hemispheric specialization of TPJ in attention, social cognition, and language functions. In general, functional specialization of the TPJ areas was stronger in the left hemisphere. The findings from the task activation correlation were highly consistent with those of the meta-analysis. Notably, there were significant differences in social cognition relevant to the three TPJ areas between HCP and CHCP datasets. Furthermore, the correlation analysis of multimodal brain maps illustrated a close relationship between the HFA map and multimodal brain maps. The consistency of maps derived in distinct analyses demonstrated the feasibility of HFA in further understanding psychological and behavioral mechanisms on neural lateralization from the perspective of hemispheric functional integration and specialization. In summary, the proposed HFA framework provides a reliable and valid functional brain mapping tool, with broad applicability in population neuroscience.

---

Homotopic brain regions refer to anatomically corresponding areas located at mirror-symmetric positions across the two cerebral hemispheres, which originate from evolutionary and developmental processes [1]. The corpus callosum provides direct structural connections that enable information integration between homotopic regions, resulting in high interhemispheric functional similarity, known as functional homotopy [2, 3]. Meanwhile, specific connections between each homotopic region and other functional networks support the lateralization of brain function, a key feature in cognitive evolution [4–6]. Functional homotopy is stronger in primary sensory areas due to higher demands for interhemispheric coordination, whereas higher-order cognitive functions exhibit more pronounced functional asymmetry, leading to reduced homotopy [7–9]. Abnormal functional homotopy has been observed in various cognitive and psychiatric disorders, suggesting its potential as a neurobiological marker for brain health [10, 11].

Homotopic functional connectivity, derived from resting-state fMRI data, quantifies interhemispheric functional coordination by directly computing the Pearson’s correlation between time series of homotopic brain regions [12]. This measure has shown substantial reliability [13, 14] and has provided initial evidence regarding interhemispheric coordination, hemispheric lateralization, and their associations with cognition and mental health [15–17]. However, this data-driven method lacks a theoretical foundation grounded in brain connectivity principles. The dual-factor generative model of the connectome [18], supported by extensive empirical research across species and developmental stages, suggests that the organization of brain connectivity is primarily shaped by two factors: (1) the spatial distance between brain regions (i.e., geometric properties), and (2) the similarity of their connectivity patterns (i.e., topological properties) [19, 20]. Traditional measures of homotopic functional connectivity primarily capture the geometric symmetry between homotopic regions, but fail to account for the similarity in their network-level topological patterns. Consequently, they are limited in accurately characterizing the full profile of interhemispheric functional similarity, which restricts both the quantification and interpretation of functional homotopy.

Recent large-scale studies of human brain networks have demonstrated the significance of connectivity profile similarity in systems neuroscience. Yeo and colleagues revealed the macroscale organization of the human resting-state connectome and proposed a cortical parcellation scheme based on intrinsic functional homogeneity [21–23]. Building on this, a homotopic functional parcellation was developed to provide prior information about the functional correspondence between homotopic regions [24]. Finn and colleagues introduced the concept of functional connectome fingerprinting, defining each brain region by its unique pattern of connectivity and enabling cross-individual and cross-regional comparisons of connectivity profiles [25, 26]. Subsequently, Margulies and colleagues defined the similarity between connectivity fingerprints as functional affinity. By applying dimensionality reduction to the whole-brain affinity matrix—that is, the matrix reflecting connectivity profile similarity between all brain regions— they identified the intrinsic axes of functional connectome organization, known as functional connectivity gradients [27]. Further research has shown that these gradients are associated with a range of neurobiological phenotypes, including brain evolution, development, behavior, and psychopathology [28, 29].

Drawing on the dual-factor generative model of the connectome, the present study introduces a new framework to quantify Homotopic Functional Affinity (HFA), defined as the similarity between the connectivity fingerprints of homotopic regions. Using resting-state fMRI datasets with high temporal (700 ms) and spatial (2 mm) resolution from the Chinese and U.S. Human Connectome Projects, we constructed whole-brain HFA maps. Test–retest reliability was assessed via linear mixed-effects modeling, demonstrating the method’s robustness for measuring individual differences. Finally, the validity of HFA was supported through cognitive association analyses in regions of interest and multimodal whole-brain map comparisons. This framework advances the methodological repertoire for studying functional homotopy and offers a more reliable and informative neuroimaging marker of individual variability in brain function.

## Methods & Materials

### Participants

#### Human Connectome Project (HCP) Dataset

The study utilized resting-state fMRI data and task-based fMRI data (during language processing and social cognition tasks) of 339 unrelated subjects (male/female = 157/182, age = 28.64 ± 3.70 years) from the HCP dataset [30]. The data were acquired on a customized 3T Siemens Skyra Connectome MRI scanner, with the following parameters: repetition time of 720 ms, echo time of 33.1 ms, 72 slices, a flip angle of 52 degrees, an in-plane field of view of 208 × 180 mm^2^, and an isotropic voxel size of 2 mm^3^. Each subject underwent four resting-state fMRI scans over two days (two scans per day with phase encoding directions from left to right and right to left). A single resting-state fMRI scan included 1200 time points, lasting 14.4 minutes. The cognitive behavioral paradigms for the tasks are described in 1.2, with two scans in opposing phase encoding directions (left to right and right to left) for each task. The single scan duration for the language processing task was approximately 3.8 minutes, containing 316 time points. The social cognition task lasted about 3.3 minutes per scan, containing 274 time points [31].

#### Chinese Human Connectome Project (CHCP) Dataset

The CHCP dataset, designed as the Chinese counterpart to the HCP dataset, is a multimodal imaging large dataset based on the Chinese population [32]. The current study included resting-state fMRI data from 217 subjects (male/female = 109/108, age = 22.37 ± 2.88 years) in the CHCP dataset, of which 185 subjects had task-based fMRI data (male/female = 90/95, age = 22.57 ± 2.99 years). The data collection process was similar to HCP, using a 3T Siemens Prisma MRI scanner. Each scan had a repetition time of 710 ms, echo time of 30 ms, and an in-plane field of view of 212 mm × 212 mm. The number of slices, flip angle, and voxel size were the same as the HCP dataset. Resting-state fMRI scans were conducted over two days, with two scans each day in opposite phase encoding directions (anterior-to-posterior and posterior-to-anterior). Each scan included 634 time points, lasting 7.5 minutes. The behavioral paradigms for language processing and social cognition tasks were consistent with HCP but were adapted into Chinese. Each task included two fMRI scans in opposing directions. The scan duration was nearly identical to HCP, with 321 time points for the language processing task and 278 time points for the social cognition task.

### Cognitive Behavioral Paradigm

#### Language Processing Task

The language processing task was designed to assess phonological and semantic processing and consisted of two runs (i.e., two separate scans). Each run included four blocks of a story comprehension task and four blocks of a math task, presented in alternating order. Each block lasted approximately 30 s. In each story block, participants listened to 5–9 sentences adapted from Aesop’s fables and were asked to choose the main theme of the story from two given options. In each math block, participants heard spoken arithmetic problems and were asked to perform addition or subtraction and select the correct answer from two choices [33]. The auditory and speech input of the math task was matched to that of the story task in terms of presentation format and attentional demands, but it did not involve semantic processing, thereby serving as a control condition for the story task.

#### Social Cognition Task

The social cognition task, adapted from the studies of Castelli et al. [34] and Wheatley et al. [35], assessed individual’s theory of mind levels through video during two rounds of scanning. Each round included five sets of geometric video tasks (2 sets of social interaction videos and 3 sets of random videos or vice versa) and five sets (each 15s) of fixed gaze tasks. In each geometric video set, subjects watched a 20s video involving shapes like squares, circles, and triangles. Subsequently, subjects had to decide whether there was social interaction among the movements of these shapes in the video, with three options: present (the movements of the shapes appeared to consider each other’s feelings and thoughts), uncertain, and absent (no interaction, movements were random).

#### MRI Data Preprocessing

The MRI data used in this study had been preprocessed by the HCP and CHCP teams using their standardized processes and were publicly shared [32, 36]. Both resting-state and task-based fMRI data preprocessing were completed using the HCP minimal preprocessing pipelines [36]. This included correction for field inhomogeneities and head motion, as well as establishing mappings between individual space and standard space. Data were then mapped to the standard 32k grayordinate space, followed by Gaussian smoothing with a Full Width at Half Maximum (FWHM) of 2mm. Resting-state fMRI data were high-pass filtered with a threshold of 2000s to remove slow drifts in the time series and underwent independent component analysis-based denoising. Task-based fMRI data were processed for individual task activations, including high-pass filtering with a threshold of 200s (for more information on data preprocessing and dataset comparisons, refer to references [31, 32, 36]).

#### Concatenation of Resting-State fMRI Time Series

To reduce random noise in the resting-state fMRI data and improve computational efficiency, a MELODIC’s Incremental Group-PCA (MIGP) method was used for concatenation processing of the time series of each vertex on the cortex after preprocessing [37]. The total length of time series for each vertex was slightly shorter than the total number of time points of an individual subject’s scan. Prior to the computation of group-level maps, all resting-state fMRI data from all participants were concatenated using this method. For the test–retest reliability analysis, the two resting-state runs acquired on the same day for each participant were first concatenated, resulting in two time series per participant (one per scan day), which were treated as repeated measurements. Additionally, all four resting-state runs from each participant were concatenated to generate an average time series, which was used in subsequent analyses assessing the association between HFA and task-evoked cognitive activation.

#### Calculation of HFA Maps

The HCP grayordinate space provides a spatially symmetric cortical template for homotopic analysis. Cortical surface vertices with identical indices in the left and right hemispheres are defined as homotopic vertex pairs. As illustrated in Figure 1, we first computed the whole-brain functional connectivity profile (i.e., connectivity fingerprint) for each homotopic vertex, reflecting its temporal synchrony (positive connectivity) and asynchrony (negative connectivity) with all other cortical vertices. The HFA at each location was then quantified as the cosine similarity between the connectivity fingerprints of the homotopic pair. A higher affinity value indicates greater functional homogeneity between the homotopic vertices, while a lower value indicates greater heterogeneity. Importantly, when computing cosine similarity, both intrahemispheric and interhemispheric connectivity of one vertex were aligned with the corresponding connectivity of its homotopic counterpart. Using the concatenated resting-state fMRI time series, we computed HFA maps at both the group and individual levels for the HCP and CHCP datasets. These included: (1) group-level maps, (2) individual test–retest maps based on data from two separate days, and (3) individual maps generated from all four concatenated runs. To enhance spatial smoothness and signal-to-noise ratio, all individual-level homotopic functional affinity maps were surface-smoothed using a Gaussian kernel with a FWHM of 5 mm.

**Figure 1.**
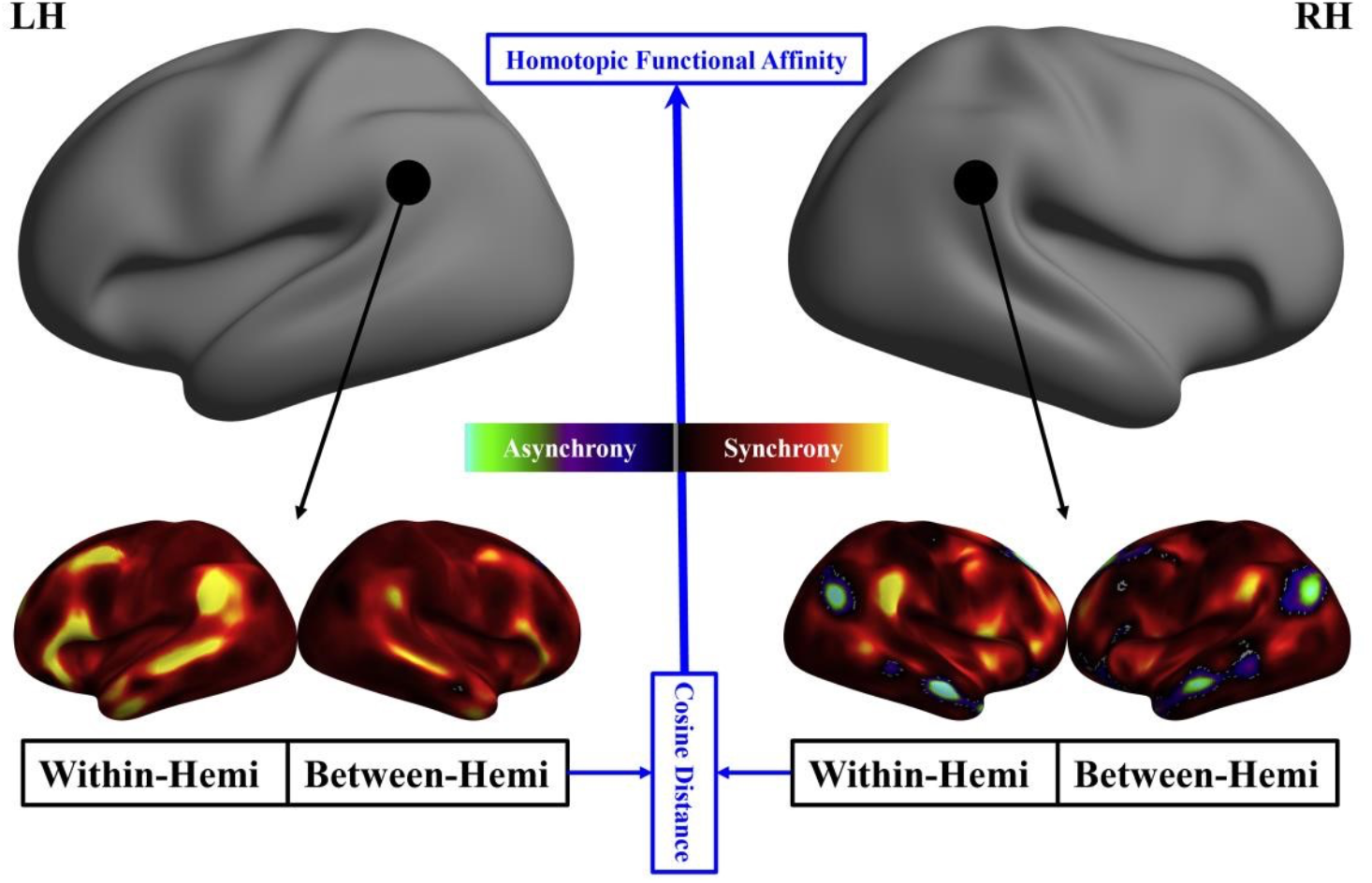
Homotopic functional affinity calculation. Given a pair of homotopic vertices on the cerebral cortex (black circled vertex in the left and right hemispheres with opposite horizontal coordinates), their whole-brain functional connectivity patterns (functional fingerprints) were calculated respectively to quantify their temporal synchrony (positive functional connectivity) and asynchrony (negative functional connectivity) with other vertices. The connectivity fingerprints of homotopic vertices were then concatenated in the order of intra-hemispheric and inter-hemispheric connectivity (as shown in the figure) respectively, and the cosine distance between the concatenated fingerprints was calculated as the homotopic affinity of the vertex.

#### Test-Retest Reliability Assessment

The test-retest reliability of the HFA measurement was assessed using a linear mixed-effects model [38-40]. Firstly, age and gender were considered as covariates to assess the within-subject variability and between-subject variability (1).

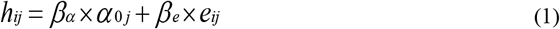

*h*_*ij*_ represents the homotopic affinity measurement value for subject *j* during measurement *i*, determined by α _0_ *j* and the model residual *e*_*ij*_. Here, 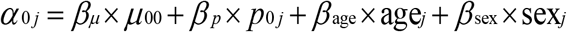. μ_00_ as a fixed effect, represents the average of all subjects’ test and retest measurements, and age_*j*_ and sex_*j*_ are the age and gender of subject *j, p*_0 *j*_ is the average of different measurements for subject *j. α* _0 *j*_ varies only with different subjects *j*. After regressing for age and gender, the between-subject variability *V*_*b*_ can be represented by the variability of *p*_0 *j*_. When subject *j* is fixed, the model residual *e*_*ij*_ comes from the variability between different measurements *i* for subject *j*. Hence, the within-subject variability *V*_*w*_ can be represented by the variability of *e*_*ij*_. By calculating the ratio of between-subject variability to total variability (sum of within-subject and between-subject variability), the Intraclass Correlation Coefficient (ICC) (2) is obtained, serving as the standard for measuring test-retest reliability.

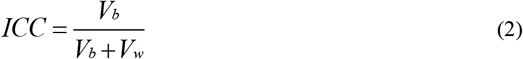

The ICC of the global mean *μ* and standard deviation *σ* of homotopic affinity, as well as the ICC for each vertex, were calculated. The calculation of the ICC on vertices required individual-level standardization of homotopic affinity values (3). For vertex *k*, its standardized homotopic affinity value *h*_*kz*_ is equal to the original homotopic affinity value *h*_*k*_ minus the individual HFA map’s global mean *μ*, divided by the global standard deviation *σ*.

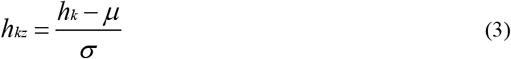

#### Validity Analysis

Based on the brain-wide distribution patterns of HFA and the test-retest reliability levels of individual differences at each vertex in different datasets, the Temporo-Parietal Junction (TPJ) - an area with high reliability and marked regional differentiation (low affinity) and sample variability - was identified as ROI for further analysis of the HFA maps. Using the HFA map from HCP, affinity edge detection near TPJ was performed to locate its subregions and extract their whole-brain functional connectivity fingerprints across different datasets, deciphering the source of their functional homotopy differentiation. Meta-analysis based on the Brainmap database (https://www.brainmap.org) [41-43] was conducted to analyze cognitive-behavioral function associations of these TPJ subregions in both hemispheres. Finally, using the method described in (3), individual HFA maps were standardized. The mean normalized HFA within each ROI and the corresponding cognitive task activation levels were extracted. Pearson’s correlation was computed between these measures, followed by multiple comparison correction. The results were then compared with findings from prior meta-analyses to evaluate their consistency.

Using Neuromaps [44], we extracted human brain gene expression maps [45], cortical evolutionary expansion maps [46], cross-species functional homology index maps [47], cortical myelination maps [48, 49], functional connectivity principal gradient maps [27], and cognitive association principal maps [50]. All maps were transformed into the standard 32k grayordinate space. We then modeled and analyzed the spatial correspondence between the averaged bilateral maps and the HFA map.

## Results

### High-Resolution Whole-Brain HFA Maps

The HFA maps exhibited highly consistent spatial distribution patterns across both the HCP dataset (Figure 2a) and the CHCP dataset (Figure 2b). Specifically, HFA across homotopic brain regions showed a gradual decline along the primary-to-association cortex gradient: the visual, somatomotor, and auditory networks demonstrated the highest HFA, followed by intermediate levels in the Ventral and Dorsal Attention Networks (VAN, DAN), and the lowest HFA in the Default Mode Network (DAN) and Fronto-Parietal Networks (FPN).

**Figure 2.**
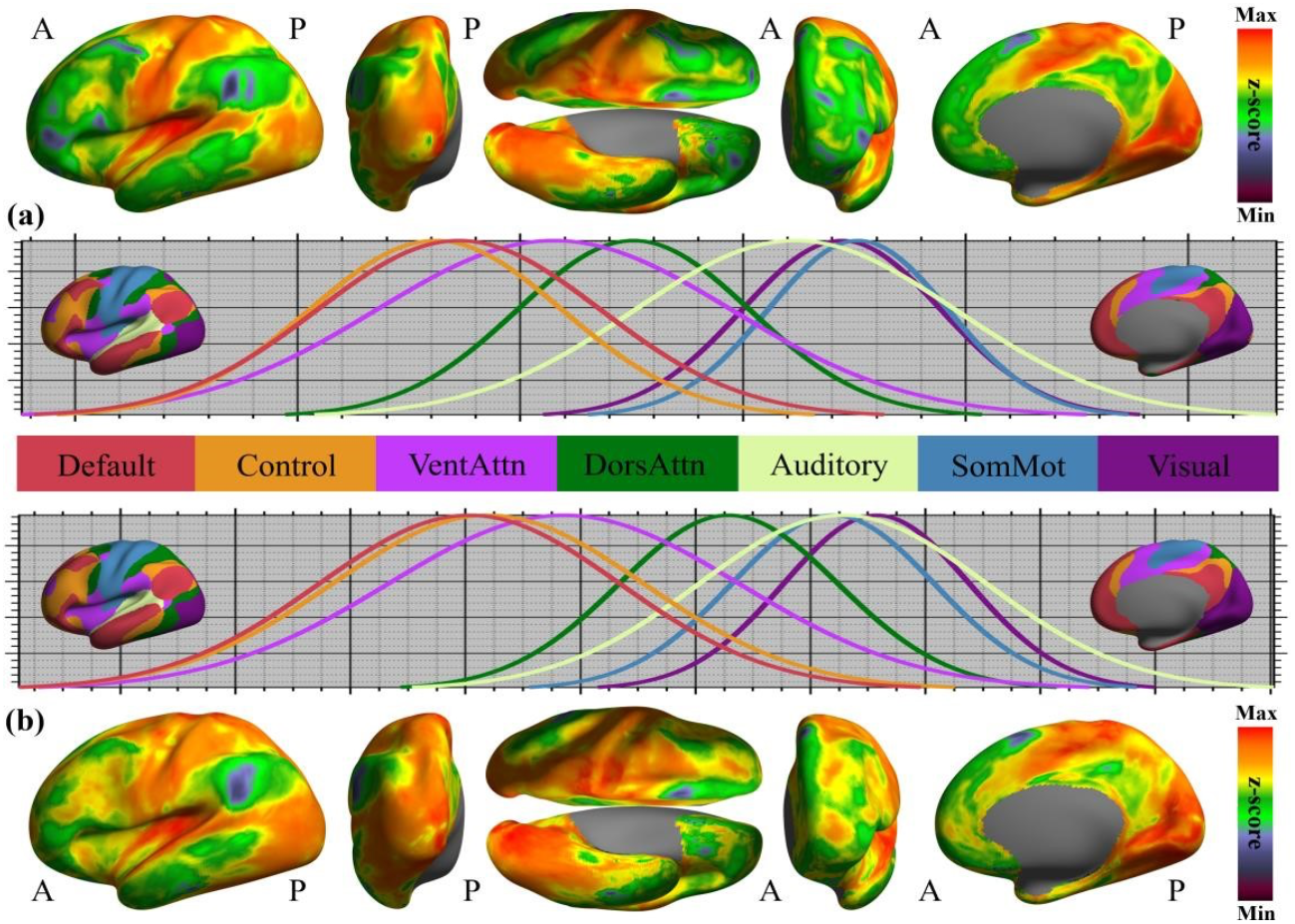
Group-level HFA of human cortex. The HFA maps of both HCP (a) and CHCP (b) were derived and projected onto the hemispheric cortical surface. For the seven large-scale functional connectivity networks in the human brain [32], a series of Gaussian models were used to fit their homotopic affinity distributions across cortex.

### Test-Retest Reliability of Individual Differences in HFA

The reliability of individual differences in HFA was quantified using ICC, with values ranging from 0 to 1. Based on 0.2 intervals, ICC was categorized into five levels: slight, fair, moderate, substantial, and near-perfect [51]. The whole-brain mean and standard deviation of HFA showed moderate reliability (HCP: ICC = 0.59 and 0.56; CHCP: ICC = 0.50 and 0.39). Vertex-wise ICC maps are presented in Figure 3. In the HCP dataset, 73.58% of cortical vertices exhibited moderate or higher reliability, compared to 53.56% in the CHCP dataset, which may reflect differences in measurement design—most notably scan duration—between the two datasets. Nevertheless, the spatial patterns of ICC across the brain were highly consistent across datasets. Specifically, HFA in association cortices showed higher test–retest reliability than in primary sensory cortices. Regions with known low BOLD signal-to-noise ratio, such as the central sulcus, orbitofrontal cortex, and insula, exhibited lower ICC values overall, although these values were consistently higher in HCP than in CHCP.

**Figure 3.**
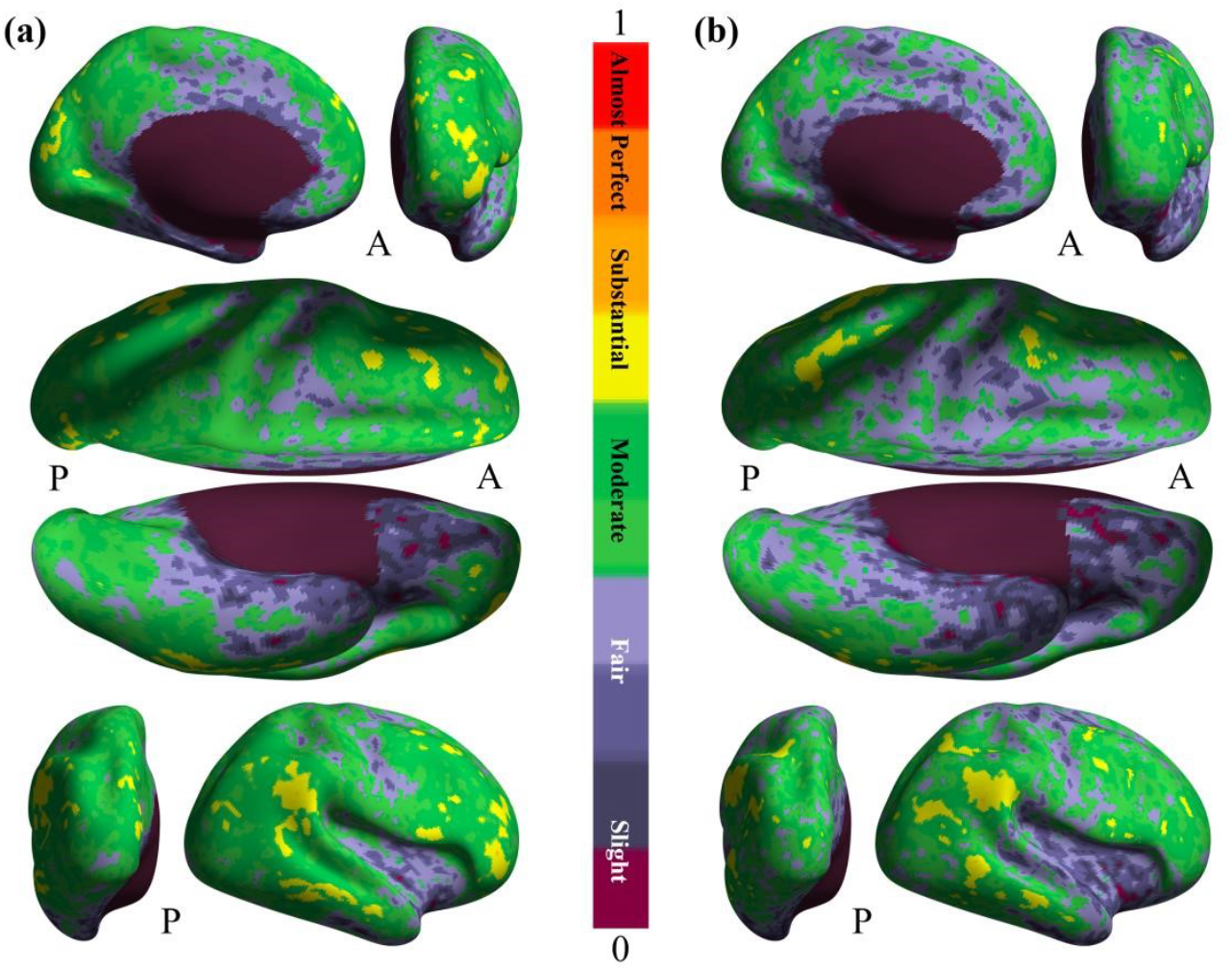
Test-retest reliability maps of measuring individual differences in HFA. Linear mixed models are employed to model individual differences in cortical HFA and calculate its test-retest reliability quantified by intraclass correlation coefficients, which are rendered onto the cortical surfaces for (a)HCP and (b) CHCP, respectively.

### Validity of Individual Difference Measurements in HFA

To evaluate the validity of the HFA map for capturing individual differences, analyses were conducted at both regional and global levels. At the regional scale, we focused on the temporoparietal junction (TPJ), a region exhibiting low HFA, by comparing its subregional differentiation and corresponding whole-brain functional connectivity fingerprints, and further validating their distinct cognitive activations. At the global scale, we assessed convergent validity by correlating the HFA map with maps of evolutionary expansion, gene expression, cortical myelination, and functional organization.

### TPJ Subregion Localization and Validation

Using an edge detection algorithm on the group-level HFA map from the HCP dataset, three subregions were identified within the TPJ: the anterior TPJ (TPJa), central TPJ (TPJc), and posterior TPJ (TPJp). The spatial boundaries of these subregions are shown in the central panel of Figure 4, with their left and right hemisphere-specific whole-brain functional connectivity fingerprints depicted on either side (HCP dataset). Specifically, the left TPJa (lTPJa) showed predominantly synchronous functional connectivity patterns, characterized by strong connectivity among the DMN, FPN, and auditory network. In contrast, its right-hemisphere homotopic counterpart (rTPJa) exhibited strongest synchrony with the VAN and strongest asynchrony with the DMN.

**Figure 4.**
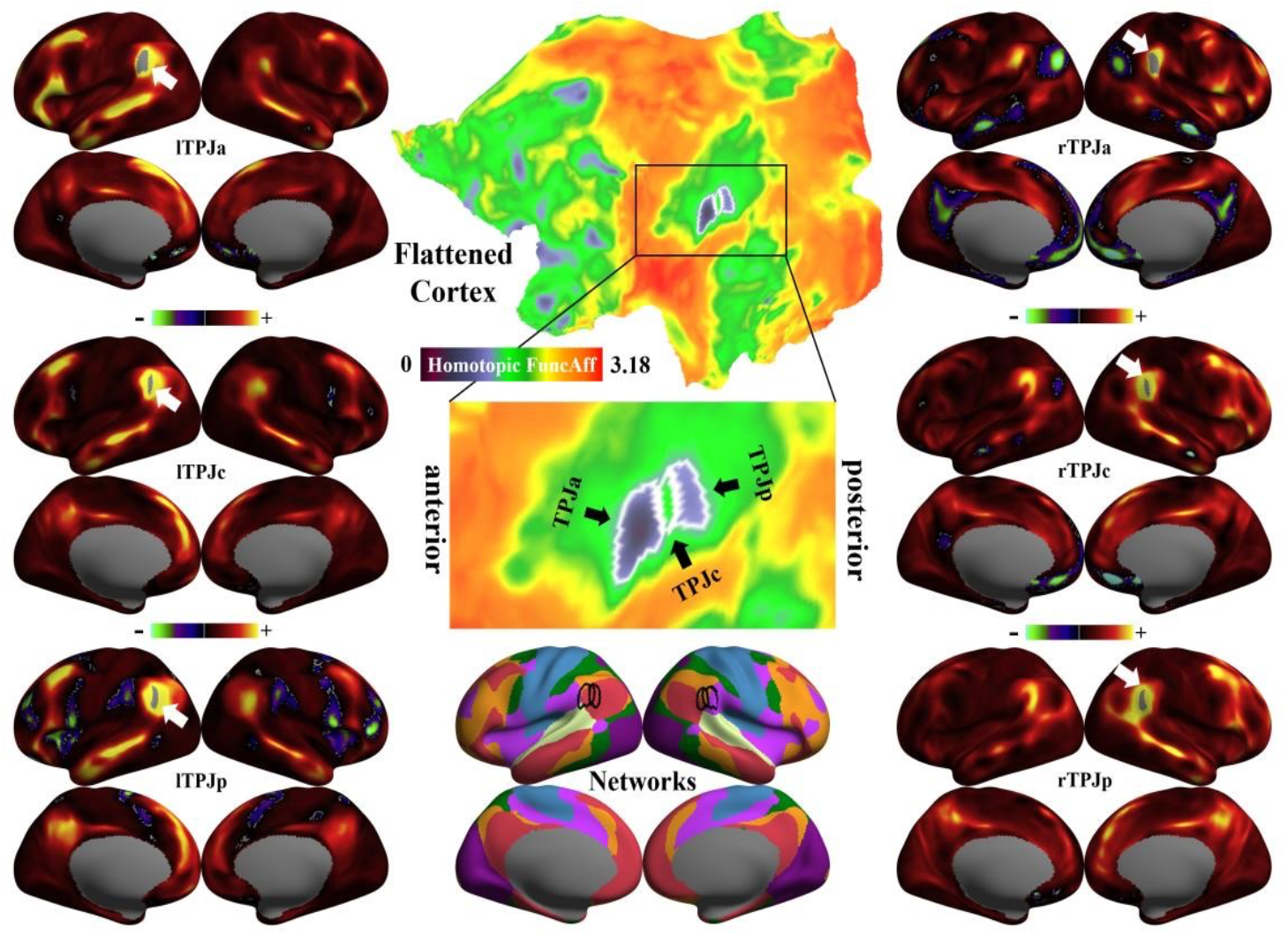
Homotopic affinities and functional connectivity fingerprints of the TPJ subregions. Based on the HFA map derived from HCP, we used an edge detection algorithm to locate the three TPJ subregions: anterior TPJ (TPJa), central TPJ (TPJc), and posterior TPJ (TPJp). The central panel shows the HFA map, the boundaries of the three TPJ subregions, and their positions within the seven large-scale functional networks [32]. The left panel demonstrates the whole-brain functional connectivity fingerprints of the left TPJ subregions, and the right panel is for their homotopic positions.

The left TPJp (lTPJp) displayed strong synchrony with the DMN and FPN, along with strong asynchrony with the VAN. Its homotopic counterpart (rTPJp) also showed strong synchrony with the DMN and FPN, but only limited asynchrony with the DMN. The central subregion TPJc showed transitional connectivity fingerprints between TPJa and TPJp. The left TPJc (lTPJc) resembled lTPJa in its synchronous connectivity pattern but lacked asynchronous patterns; rTPJc was similar to rTPJp in synchrony and partially overlapped with rTPJa in asynchrony.

In the CHCP dataset, whole-brain functional connectivity fingerprints of bilateral TPJa and TPJp closely matched those in the HCP dataset. However, TPJc connectivity patterns differed markedly: lTPJc more closely resembled lTPJp, while rTPJc was more similar to rTPJa. Both lTPJc and rTPJc exhibited asynchronous connectivity, reducing the similarity between the two and making it difficult to distinguish TPJc from TPJa and TPJp based solely on HFA, despite their similarly low HFA values.

The results of the meta-analytic decoding for the six TPJ subregions (three per hemisphere) are shown in Figure 5. A comparison of the cognitive word clouds across hemispheres revealed that functional specificity was generally greater in the left TPJ, while the right TPJ subregions were associated with more overlapping cognitive processes. Similar to the whole-brain functional connectivity fingerprints, the associated cognitive functions of these subregions displayed transitional patterns along the TPJ axis.

**Figure 5.**
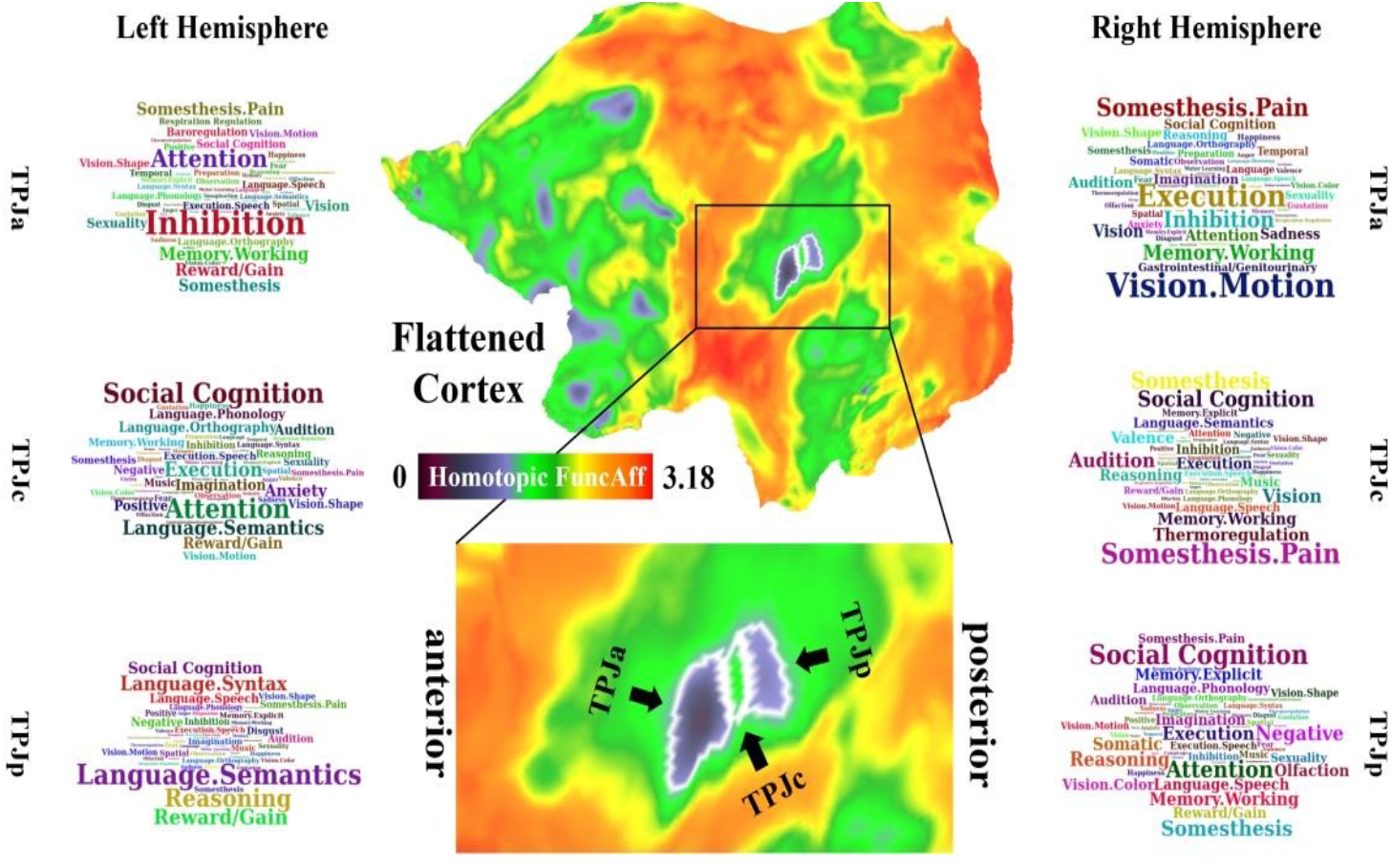
Word cloud maps of cognitive association with the three TPJ subregions. The central panel depicts the human brain functional affinity map and the boundaries of the three TPJ subregions. The left panel demonstrates the word clouds derived from the meta-analysis on cognitive function of the left TPJ subregions, and the right panel is for their homotopic positions.

TPJa was most strongly associated with inhibition, attention, and executive control, with additional associations to somatosensory processing observed in the right hemisphere. TPJc was primarily linked to social cognition, and its right-hemisphere counterpart also showed relevance to somatosensory processes. TPJp was most strongly associated with language-related functions, particularly in the left hemisphere, whereas the right TPJp was more closely related to social cognition.

Building on the meta-analytic profiles, we further examined the relationship between HFA and task activation during language comprehension and social cognition tasks for each of the six TPJ subregions (Table 1). For the HCP dataset, in the language task, the Story-Math contrast revealed significant and opposite correlations between HFA and activation in lTPJa and rTPJp. A similar pattern of opposite correlations was also observed for bilateral TPJc, corresponding to the direction of effects seen in lTPJa and rTPJp. These relationships were also evident in the Story condition alone, while no significant correlations were found under the Math condition. In the social cognition task, significant negative correlations between HFA and activation were found for both rTPJc and rTPJp under the Social Interaction and Random conditions.

**Table 1.**
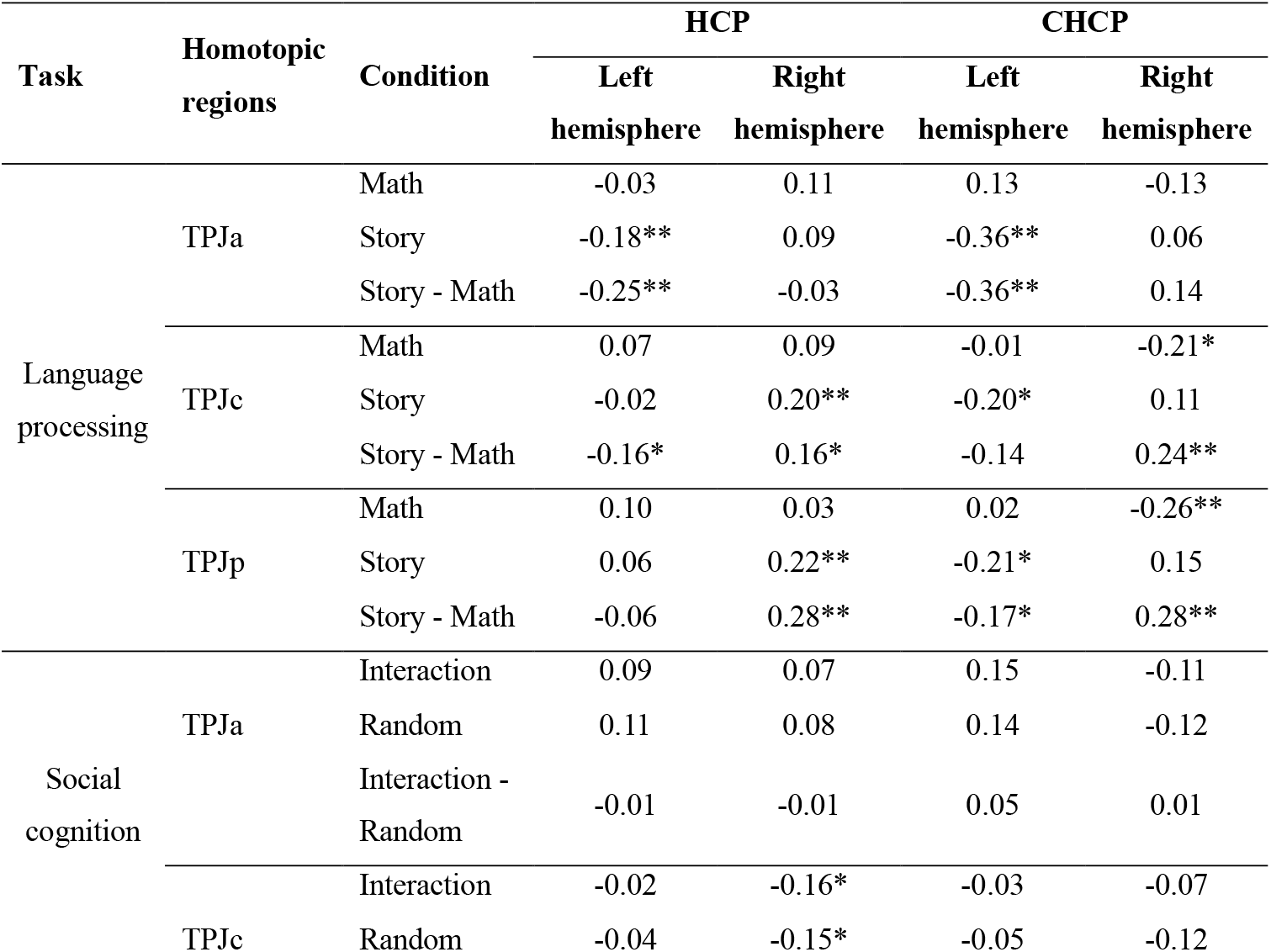
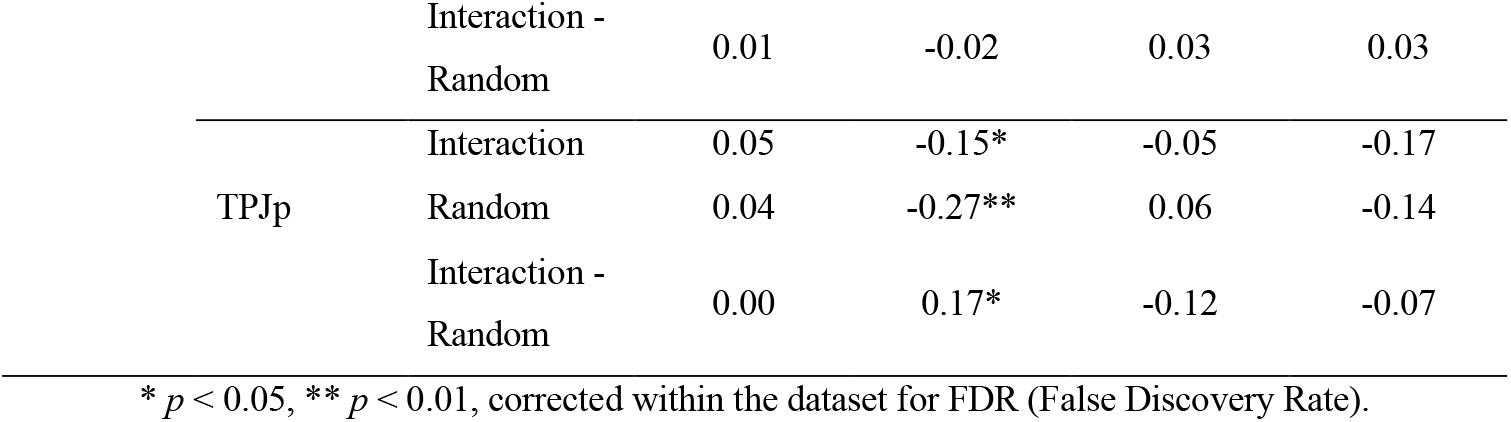
Association of homotopic functional affinity in temporo-parietal junction subregions with cognitive task activation.

In the CHCP dataset, the pattern of HFA–activation correlations observed during the Story-Math contrast was consistent with the HCP findings. In the Story condition, all three left TPJ subregions exhibited significant negative correlations between HFA and activation. Under the Math condition, significant correlations were also observed for rTPJc and rTPJp. For the social cognition task, the CHCP results showed similar trends to those in HCP, though none of the correlations reached statistical significance.

### Multimodal Associations of the HFA

The cortical HFA map showed significant correlations with all six reference brain maps included in the analysis (Figure 6 presents results from the HCP dataset; corresponding spatial association maps from CHCP are provided in the Supplementary Materials). Among these, the strongest association was observed with the principal gradient of functional connectivity (HCP: r = −0.77; CHCP: r = −0.73). The negative correlation indicates that regions with higher HFA tend to occupy lower positions in the functional hierarchy, whereas regions with lower HFA tend to occupy higher-order positions along the gradient. Cortical evolutionary expansion was also negatively correlated with HFA (HCP: r = −0.53; CHCP: r = −0.45), suggesting that regions undergoing greater expansion during evolution typically exhibit lower HFA, whereas more conserved regions tend to show higher HFA. In contrast, HFA was positively correlated with three other maps—gene expression similarity, cross-species functional homology index, and cortical myelination—with correlation coefficients ranging between 0.53 and 0.60. This indicates that regions with higher HFA tend to exhibit stronger genetic constraints, greater cross-species functional similarity, and more pronounced myelination. Conversely, regions with lower HFA are characterized by reduced constraints in these domains. Finally, HFA showed the weakest association with the Neurosynth-based cognitive activation synthesis map (HCP: r = 0.35; CHCP: r = 0.38), implying that regions with higher HFA tend to have more consistent cognitive associations, while regions with lower HFA may require further investigation to elucidate their cognitive roles.

**Figure 6.**
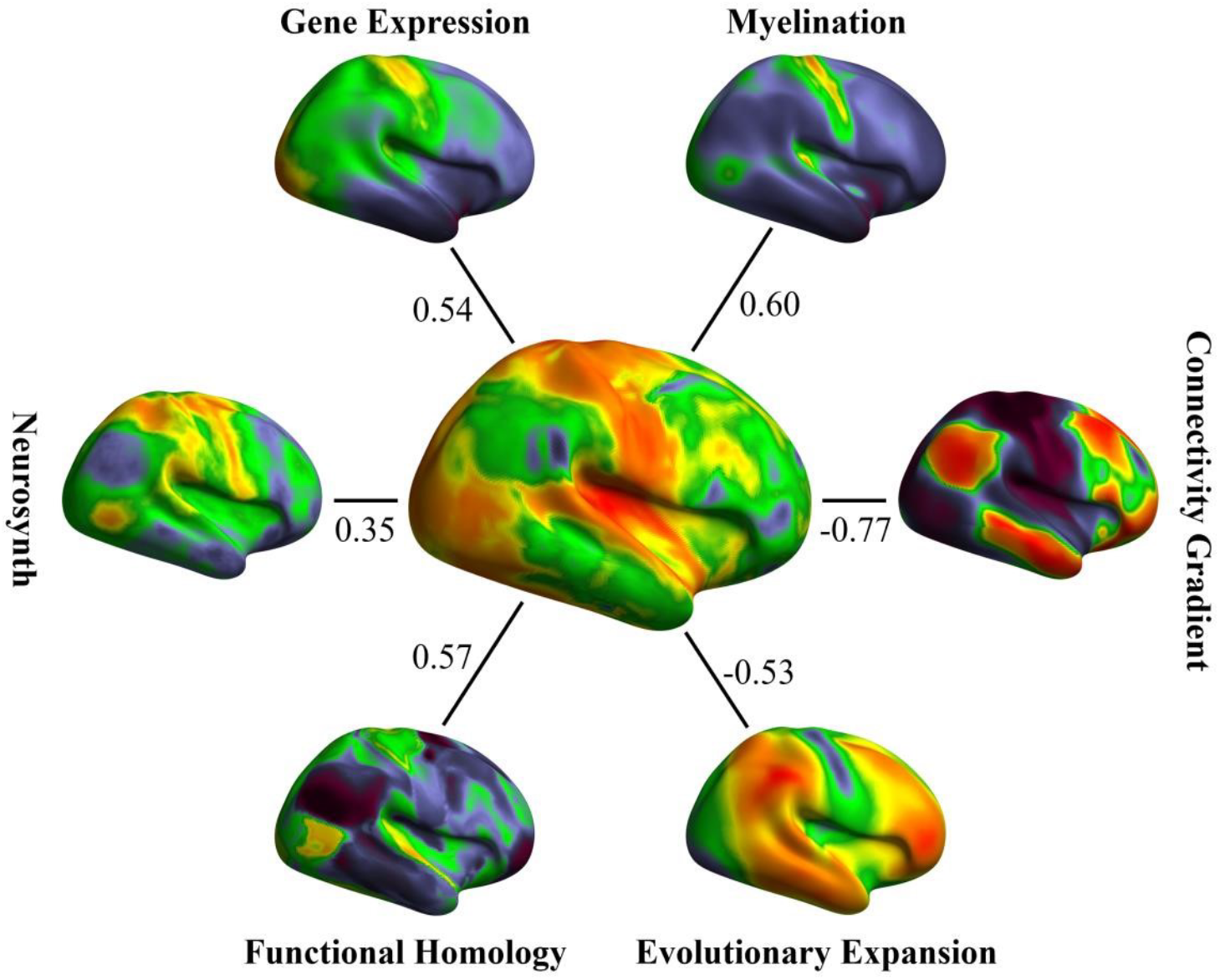
Multimodal map association of HCP HFA map.

## Discussion

This study introduces a novel metric—HFA—and systematically validates its reliability and validity in measuring individual differences. HFA serves as a robust indicator of interhemispheric functional integration and regional specificity, with unique relevance to genetic, evolutionary, and organizational features of the human brain. Using this framework, we revealed the functional complexity of the TPJ, demonstrating the method’s scientific value and potential applications in regional functional differentiation, resting–task state correspondence, and the investigation of cultural and ethnic differences in brain function.

### Reliability of the HFA Method

Despite the widespread application of functional magnetic resonance imaging (fMRI) in brain function studies, the reproducibility of related findings remains challenging [52, 53]. One primary reason for this low reproducibility is the insufficient reliability of selected indices in experimental designs [54]. The reliability of the global mean homotopic functional affinity atlas exceeds 0.5, with vertex-wise homotopic functional affinity generally surpassing 0.4. This reliability outperforms general resting-state functional connectivity and traditional task activation methods, approaching other highly reliable resting-state fMRI metrics [13, 14], thus establishing it as a dependable measure of brain functional homotopic integration and lateralization.

In associative cortical areas like the DMN and FPN, the reliability of HFA is generally higher and consistent across datasets, aligning with the distribution patterns of other resting-state fMRI reliability metrics [14, 55]. These cortical areas exhibit significant inter-individual variability [56] but relatively smaller intra-individual variation [57, 58], making it easier to distinguish between individuals based on their activity patterns [59]. In regions like the central sulcus, orbitofrontal cortex, and insula, where blood oxygen level-dependent signals are constrained, the reliability of homotopic functional affinity in the HCP dataset tends to be higher than in the CHCP. This discrepancy may stem from differences in scanning duration across datasets, with HCP’s resting-state fMRI scanning duration being twice that of CHCP. Longer scanning durations are known to provide more stable time series, reducing the impact of random fluctuations and enhancing test-retest reliability [60, 61].

### Multimodal Biological Correlates of HFA

The HFA map reveals a gradual enhancement of functional specificity along the primary-associative cortical gradient axis. This axis dominates different dimensions of human brain architecture and function [28, 62]. Our study corroborates the decisive role of this gradient axis in the intrinsic functional representation of the human brain, from the perspective of functional integration and specialization in homotopic regions. Further, multimodal brain maps’ correlation analysis unveils the potential genetic, evolutionary, and cognitive significance underlying this spatial distribution pattern.

Hemispheric asymmetry in gene expression emerges during early embryonic development [63]. In lateralized networks such as the language system, hemisphere-specific expression of genes related to electrophysiological and neurotransmitter pathways has also been documented [64]. Evolutionarily, primary cortices emerged earlier and serve foundational sensory and perceptual roles. The HFA framework assumes symmetric homotopy as a starting point in evolution. With cortical expansion, especially in association cortices and higher-order cognition [65], regions showing the most pronounced expansion also exhibit the strongest functional lateralization [4]. Language-and control-related areas show greater asymmetry in humans than macaques [6], and display lower cross-species functional homology [47]. These findings emphasize the close link between cognitive evolution and hemispheric specialization [8], supporting the theoretical validity of the HFA model and offering insights into the biological basis of lateralization.

### Functional Specificity of TPJ

The TPJ is involved in multiple high-order cognitive processes and occupies a unique position in the human brain [66]. No homologous region with comparable function has been identified in the macaque brain, particularly within the inferior parietal lobule [47]. The cortical HFA map captured the TPJ’s functional specialization, revealing fine-grained inter-subregional and interhemispheric differentiation. Functionally, TPJ subregions transition from ventral executive control to dorsal social cognition and then to posterior language processing [67]. In the right hemisphere, rTPJa and rTPJc are components of the ventral attention and FPN, respectively. Meta-analytic results revealed their associations with somatic pain and visual processing, pointing to a tight coupling between somatosensory input, attentional orientation, and executive control [68, 69]. Rajimehr et al. found that left-hemisphere activations during language tasks correspond to right-hemisphere activations during social tasks within core TPJ regions [70]. Our findings replicate this hemispheric complementarity in TPJp, offering valuable insight into the neural basis of language–social cognition interaction and related disorders. Furthermore, the directionality of correlations between HFA and task activation across TPJ subregions aligns with resting-state connectivity fingerprint patterns, highlighting the potential of TPJ connectivity architecture in predicting intra- and interhemispheric language information flow.

### Cultural and Ethnic Diversity in the HFA Map

This study generated cortical HFA maps based on two high-resolution datasets from American (HCP) and Chinese (CHCP) populations. While the overall patterns were highly consistent, systematic differences emerged in specific regions. Notably, the relative HFA levels of TPJc versus TPJa/TPJp differed between datasets. These divergences likely reflect population differences. Cultural and genetic heterogeneity between U.S. and Chinese participants encapsulates a range of evolutionary and environmental factors contributing to cognitive diversity [71,72].

Given TPJc’s primary association with social cognition, intergroup differences in its HFA may reflect culturally shaped neural mechanisms. Supporting this, the correlation between task activation and HFA in the social cognition task was significant in HCP but absent in CHCP. Conversely, under the arithmetic condition of the language task, TPJ activation correlated significantly with HFA in CHCP but not in HCP. Cultural context is known to influence learning and reasoning styles [73,74], and our findings provide evidence for the neurobiological imprint of cultural and ethnic diversity on brain function.

### Future Directions

Grounded in the dual-factor model of human brain connectivity, this study proposed a novel metric— HFA—and demonstrated its potential for advancing emerging functional brain mapping techniques and deconstructing the cognitive mechanisms of the human mind. From a methodological perspective, HFA can be regarded as a specific subset within the broader framework of functional connectivity gradients. The current findings revealed a strong spatial similarity between the HFA map and the principal functional gradient. However, the relative contribution of HFA to gradient derivation, the nuanced distinctions between HFA and the principal gradient, and their respective cognitive implications remain to be fully elucidated. Furthermore, the current HFA computation incorporates whole-brain connectivity profiles. Given that different brain regions vary in their hemispheric versus intrahemispheric connectivity preferences, further research is warranted to disentangle the distinct roles of interhemispheric and intrahemispheric affinity in shaping HFA.

From an evolutionary and developmental perspective, our findings suggest a close relationship between HFA and human cortical evolution. It remains an open question whether similar HFA patterns exist in non-human species and whether cross-species comparisons of HFA could shed light on the evolutionary trajectory of the human brain. During development, the dual-factor generative model posits that the influence of homogeneity factors strengthens while spatial constraints weaken over time [18,19]. Since HFA was conceptually derived from this framework, future work is needed to investigate whether individual-level HFA maps follow such developmental trajectories.

Finally, this study highlights the sensitivity of HFA to sociocultural differences, particularly in the TPJ, suggesting its promise as a tool for detecting culturally and ethnically shaped patterns of brain activity. These findings also call attention to the need for cultural adaptability in experimental paradigms used in cross-cultural research. Future studies could incorporate more cross-cultural datasets to replicate and extend the current findings, and further leverage the HFA framework to uncover neurocognitive mechanisms underlying cultural and ethnic diversity.

## Declaration of Conflicting Interests

The authors declared no potential conflicts of interest with respect to the research, authorship, and/or publication of this article.

## Notes

### Competing Interest Statement

The authors have declared no competing interest.

### Summary of Updates

All main text was revised in terms of language and writing style.

